# A multi-*b*-value test-retest diffusion MRI brain dataset for model validation and reproducibility assessment

**DOI:** 10.64898/2026.08.23.746449

**Authors:** Tomasz Pieciak, Irene Guadilla, Dominika Ciupek, Rafael Navarro-González, Susana Merino-Caviedes, Pablo Villacorta-Aylagas, Laura Magdaleno Humayor, Marco Villa Aparicio, Jorge Rueda-Ramos, Raquel Santiesteban Mendo, Raúl Moro Boyero, Antonio Tristán Vega

**Affiliations:** Laboratorio de Procesado de Imagen (LPI), ETSI Telecomunicación, Universidad de Valladolid, Valladolid, Spain; Instituto de Investigación Biosanitaria de Valladolid, IBioVALL, Valladolid, Spain; Sano Centre for Computational Medicine, Kraków, Poland; Grupo de Comunicaciones Ópticas (GCO), ETSI Telecomunicación, Universidad de Valladolid, Valladolid, Spain; Laboratorio de Técnicas Instrumentales, Universidad de Valladolid, Valladolid, Spain; Hospital Clínico Universitario de Valladolid, Valladolid, Spain

**Keywords:** diffusion MRI, diffusion tensor imaging, reproducibility, data, dataset, test-retest, brain

## Abstract

Transparent assessment of diffusion magnetic resonance imaging (dMRI) techniques with empirical verification of confounding factors requires adequately designed protocols and collected datasets. Publicly available diffusion-weighted MR datasets often provide limited sampling across *b*-values, making it difficult to study optimal acquisition protocols or the relationships between different processes occurring in brain tissue. In this work, we introduce a new densely sampled longitudinal test-retest diffusion-weighted MR dataset of the brain. Our dataset was collected from eleven healthy volunteers, each scanned four times: two sessions on consecutive days, which form the test data, followed by two additional sessions completed one week later (retest data). The data were acquired using twenty-two *b*-values ranging from 10 to 3000 s/mm^2^, along with structural *T_1_*-weighted scans. Potential applications of the dataset include, but are not limited to, assessing longitudinal reproducibility and reliability of quantitative metrics, evaluating robust and outlier-resistant estimation techniques, investigating experimental factors affecting estimation procedures, and verifying optimal acquisition protocols for different signal models. The dataset is publicly available in raw and fully preprocessed variants.

## 1. Background & Summary

Diffusion magnetic resonance imaging (dMRI) is a well-established non-invasive imaging technique used to probe the diffusive motion of water molecules in biological tissues, including neural tissue (Basser et al., 1994a; Le Bihan, 2003). Over the past three decades, a broad spectrum of mathematical formulations has been developed to quantify the diffusion-weighted MR signal (see for example, Basser et al., 1994b; Jensen et al., 2005; Özarslan et al., 2012; Zhang et al., 2012; Daducci et al., 2015; Jelescu et al., 2017), enabling the investigation of brain microstructural alterations associated with neurological and psychiatric disorders (Kubicki et al., 2007; Pasternak et al., 2018), the impact of environmental factors on neurodevelopmental disorders (Lewandowska et al., 2025), as well as the characterisation of brain changes over time (Westlye et al., 2010; Cox et al., 2016; Pieciak et al., 2023; Kim et al., 2026).

Open access to diverse MRI datasets of the brain is crucial for transparent and reproducible evaluation of computational techniques used to process the data (Nichols et al., 2017; Markiewicz et al., 2021). In the context of dMRI, these include custom preprocessing pipelines, data modelling techniques for quantifying neural tissue properties, tractography algorithms for reconstructing the geometry of fibre bundles, and tractometry, which quantifies the properties of white matter tracts. Publicly available repositories, such as the Human Connectome Project (Van Essen et al., 2012; Bastiani et al., 2019), UK Biobank (Alfaro-Almagro et al., 2018), MASSIVE (Froeling et al., 2017), and MICRA (Koller et al., 2021), have greatly supported such evaluations by enabling not only extensive studies of the robustness of new algorithms but also assessments of their reproducibility, reliability, and uncertainty. In addition to the datasets mentioned above, numerous diffusion-weighted MR databases of the brain have recently been shared with the community, including raw k-space data acquired using a high-field scanner (Wang et al., 2025), connectome data obtained using a high-gradient strength system (Tian et al., 2022), and other collections enabling the evaluations of mathematical models used to represent the acquired signal in intra- and inter-scanner scenarios (Tong et al., 2020; Cai et al., 2021; Poulin et al., 2022; Warrington et al., 2025). These databases were collected using scanners from various vendors, acquisition protocols, including different voxel resolutions, diffusion weightings (represented by *b-*values), and distributions of diffusion-sensitising gradient directions. Yet, densely sampled, repeatedly acquired diffusion-weighted MR datasets of the brain remain limited in the public domain, thereby limiting comprehensive evaluations of recent diffusion-weighted MR signal representation techniques and assessments of their potential confounding factors. The availability of such densely sampled *b*-value datasets is essential not only for assessing the reproducibility of diffusion models, for which databases such as MICRA (Koller et al., 2021) or travelling subjects datasets (Tong et al., 2020; Warrington et al., 2025) were primarily designed, but also for the empirical validation of optimal acquisition protocols (Parvathaneni et al., 2018; Drenthen et al., 2023; Ciceri et al., 2024). Furthermore, not all publicly available repositories provide unprocessed data, which prevent from applying custom preprocessing pipelines for noise removal, motion artefact correction, and compensation for other artefacts, such as those arising from variations in the B0 and B1 magnetic fields.

In this work, we present an *in vivo* diffusion-weighted MR dataset acquired in a longitudinal test-retest setting across twenty-two *b*-values ranging from 10 to 3000 s/mm², along with structural *T_1_*-weighted scans. The dataset comprises acquisitions from eleven healthy volunteers, each scanned four times: two sessions on consecutive days, which form the test data, followed by two additional sessions one week later (retest data). The dataset has been shared in raw and fully preprocessed variants, enabling its immediate use with a wide range of diffusion MRI techniques, including Intravoxel Incoherent Motion (IVIM; Le Bihan, 2019) for the low-*b*-value regime, Diffusion Tensor Imaging (DTI; Basser et al., 1994b), a standard signal representation technique used in clinical settings, methods applicable to moderate diffusion weightings, such as Diffusion Kurtosis Imaging (DKI; Jensen et al., 2005), as well as moderate to higher diffusion weightings, including the Mean Apparent Propagator MRI (MAP-MRI; Özarslan et al., 2012), the Neurite Orientation Dispersion and Density Imaging (NODDI; Zhang et al., 2012) biophysical model and the Spherical Means Technique (SMT; Kaden et al., 2016). Potential applications of the presented dataset include, but are not limited to: 1) evaluating robust, outlier-resistant, or physics-informed estimation techniques (Haije et al., 2020; Tristán-Vega et al., 2023; Anania et al., 2026), 2) investigating relationships between different processes, e.g., the effect of kurtosis on the estimation of the free-water volume fraction (Pieciak et al., 2026a), 3) establishing optimal acquisition protocols for different signal representations and biophysical models (Parvathaneni et al., 2018; Drenthen et al., 2023), 4) assessing experimental factors (e.g., *b*-values and the number of diffusion gradients) affecting signal representations (Hutchinson et al., 2017; McKinnon et al., 2017), 5) developing data normalization and/or harmonisation procedures to minimize the signal variations between sessions, 6) assessing the longitudinal reproducibility and reliability of quantitative biophysical and propagator-based metrics (Lehmann et al. 2021; Boudreau et al. 2025; Pieciak et al., 2026b), tractometry (Lerma-Usabiaga et al., 2020; Taguma et al., 2026) or structural connectivity between brain regions (Girard et al., 2023). The dataset is available in the OpenNeuro repository under the following link: https://openneuro.org/datasets/ds008695.

## 2. Methods

### 2.1. Sample population

Eleven healthy volunteers (5F/6M, aged: 24–48/23–45, mean ± SD: 32.8 ± 9.2/32.2 ± 9.0) gave informed written consent to acquire and share data anonymously. The volunteers have not reported any neurological or neuropsychiatric disorders, nor any head trauma in the past. Participants were recruited through dissemination of information about the study within the university. Exclusion criteria were: head trauma, volunteers not compatible with our scanning procedure (e.g., individuals with implants, pacemakers, intrauterine devices), and pregnant volunteers. MRI acquisitions were approved by the local Ethics Committee of Hospital Clínico Universitario de Valladolid, Valladolid, Spain (PI: 14–197).

### 2.2. Data acquisition

The dataset was acquired using a Philips Achieva dStream 3T scanner (Philips, Best, Netherlands) located at the Hospital Clínico Universitario de Valladolid, Valladolid, Spain. The scanner is equipped with a 32-channel head coil. The gradient system has a maximum gradient strength of 62 mT/m and a maximum slew rate of 100 mT/m/ms. For each volunteer, the acquisition was divided into four sessions (see Fig. 1a): two sessions performed on consecutive days (test data) and two repeated sessions performed one week later also on consecutive days (retest data). For one volunteer (sub-01), due to time constraints, the test data were acquired during two sessions on the same day, whereas the retest data were acquired during two sessions on the same day one week later.

**Figure 1.**
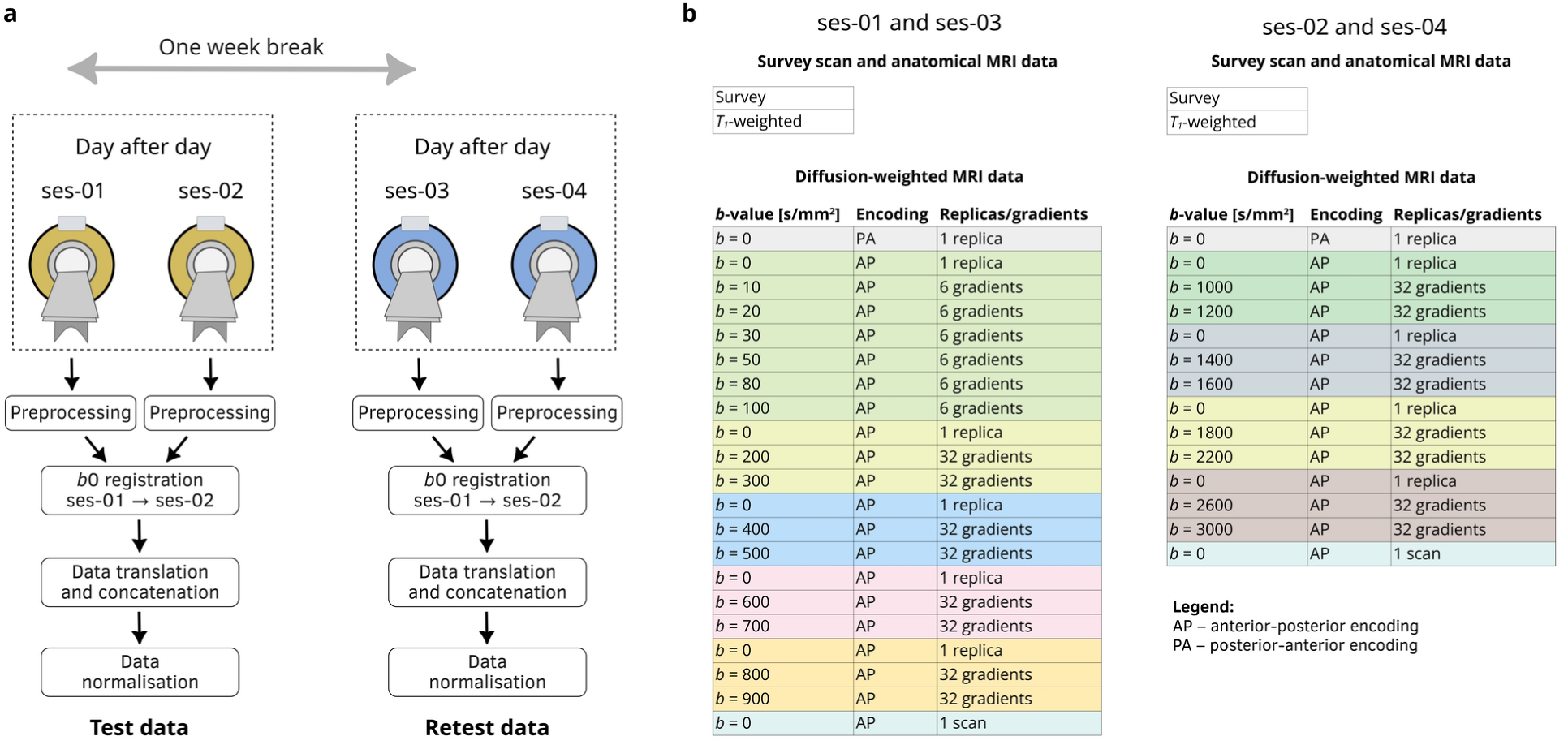
**a)** Graphical overview of data acquisition and processing procedures. Each volunteer was scanned four times on the same scanner: two sessions on consecutive days (ses-01 and ses-02), followed by two additional sessions one week later (ses-03 and ses-04). Data collected during each session were preprocessed separately, registered across sessions using *b* = 0 volumes by aligning lower *b*-value datasets to the higher *b*-value datasets, translated and concatenated, and finally normalised across volumes within each session to form the test and retest datasets **b)** Summary of the scanning protocol, including the applied diffusion-weightings represented by the *b*-values, phase-encoding directions, and the number of diffusion-sensitising gradient directions or replicas for non-diffusion-weighted volumes (*b* = 0). Each colour indicates a separate acquisition package.

#### *T_1_*-weighted MR data

Anatomical data were acquired using the 3D Turbo Field Echo (TFE) gradient-echo sequence. The acquisition setup: echo time (TE): 3.69 ms, repetition time (TR): 8.08 ms, flip angle: 8°, field of view (FOV): 240 × 240 mm^2^, acquisition voxel size: 1 × 1 × 1 mm^3^. The compressed sensing SENSE (CS-SENSE) with Cartesian sampling with an in-plane acceleration factor of 2 was used to acquire data. Reconstruction parameters: voxel size: 0.9375 × 0.9375 × 1 mm^3^, reconstruction matrix: 256 × 256 with 177 sagittal slices covering the brain.

#### Diffusion-weighted MR data

Images were acquired using a single-shot echo-planar imaging (EPI) sequence with anterior-posterior (AP) phase-encoding direction using the following parameters: TE: 95 ms, TR: 7000 ms, EPI factor: 47, effective echo spacing: 0.56 ms, flip angle: 90°, FOV: 240 × 240 mm^2^, acquisition matrix: 96 × 94, acquisition voxel size: 2.5 × 2.5 × 2.5 mm^3^. Data were acquired using Cartesian sampling with a parallel imaging acceleration factor of 2. Data were reconstructed on a 128 × 128 matrix and 55 axial slices covering the brain, which corresponds to an in-plane resolution of 1.875 × 1.875 mm^2^ and slice thickness 2.5 mm.

In total, twenty-two *b*-values were used in the acquisition procedure spread over two sessions:

- ses-01 and ses-03: *b* = 10, 20, 30, 50, 80, 100 s/mm^2^ (6 diffusion gradient directions for each shell), 200, 300, 400, 500, 600, 700, 800, 900 s/mm^2^ (32 diffusion gradient directions per shell), and six non-diffusion-weighted volumes, i.e., volumes acquired at *b* = 0 s/mm^2^,
- ses-02 and ses-04: *b* = 1000, 1200, 1400, 1600, 1800, 2200, 2600, 3000 s/mm^2^ (32 diffusion gradient directions per shell), and five non-diffusion-weighted volumes.

Diffusion-weighted MR acquisitions were organized into so-called packages. Each package, except the last one, consists of a single non-diffusion-weighted volume followed by a set of diffusion-weighted volumes (see Fig. 1b). This acquisition scheme facilitates correction of time-varying signal drift, which requires non-diffusion-weighted MR volumes to be acquired in an interlaced manner with diffusion-weighted volumes (Vos et al., 2017). In total, 298 volumes were acquired in sessions ses-01 and ses-03, and 261 volumes in ses-02 and ses-04, respectively. One additional non-diffusion-weighted volume per session was acquired with posterior-anterior (PA) phase encoding to correct for susceptibility-induced artefacts. The total acquisition time was 49 min 29 s in sessions ses-01 and ses-03, and 43 min 41 s in sessions ses-02 and ses-04.

### 2.3. Data processing

#### Data extraction

Datasets were converted from DICOM (Digital Imaging and Communications in Medicine) to NIfTI (Neuroimaging Informatics Technology Initiative) format using the dicm2nii tool v2026.03.19 (https://github.com/xiangruili/dicm2nii), which preserves the acquisition order of diffusion-weighted volumes acquired in a single package. The JSON files for the raw data were also generated using the dicm2nii tool v2026.03.19.

#### Data pseudonymisation

All imaging files were pseudonymised at two levels prior to sharing: 1) removal of sensitive meta-data from the headers of the NIfTI and JSON files using an in-house script written in the Python programming language, and 2) face masking of *T_1_*-weighted MR data using the FSL FMRIB Software Library v6 tool fsl_deface (https://fsl.fmrib.ox.ac.uk/). Subject re-identification (e.g., in the case of consent withdrawal by a volunteer) is possible only through access to separate and securely stored database that is not publicly accessible. The data were handled in accordance with Regulation (EU) 2016/679 of the European Parliament and of the Council, and Spanish Organic Law 3/2018 (LOPDGDD). These data are considered raw. Raw diffusion-weighted MR data retain the original int16 data type, whereas the face-masked *T_1_*-weighted MR data use a single-precision floating-point (float32) data type.

#### Data preprocessing

The diffusion-weighted MR data acquired at each of the four sessions were preprocessed separately using the following pipeline: 1) noise estimation and denoising using the Marčenko-Pastur Principal Component Analysis (MP-PCA) technique with a patch of size 5 × 5 × 5 voxels, implemented in the MRtrix3 dwidenoise command (Veraart et al., 2016a; Veraart et al., 2016b; Cordero-Grande et al., 2019; Tournier et al., 2019), 2) removal of Gibbs artefacts using the MRtrix3 mrdegibbs (Kellner et al., 2016), 3) estimation of susceptibility-induced distortions using the FSL FMRIB Software Library v6 topup tool (Analysis Group, FMRIB, Oxford, UK; Andersson et al., 2003; Smith et al., 2004), 4) correction for head movements and eddy current using the FSL eddy (Andersson et al., 2016), 5) correction for B1 field intensity variations using the N4 algorithm implemented in ANTs v2.6.3 (Tustison et al., 2010), and 6) time-varying drift correction (Vos et al., 2017). Noise estimation and denoising were applied separately to each package to minimise the underestimation of the noise level and consequent overestimation of the signal-to-noise ratio (SNR) resulting from differences in signal intensity between distant *b*-values. The *T_1_*-weighted MR data were not subjected to any preprocessing other than face masking.

#### Data registration between sessions

The preprocessed acquisitions with higher SNR, i.e., ses-01 and ses-03, were transformed and concatenated with their lower SNR counterparts, ses-02 and ses-04, respectively (i.e., ses-01 → ses-02 and ses-03 → ses-04), to obtain test and retest datasets (see Fig. 1a). First, the non-diffusion-weighted MR volumes were affinely registered using the FSL v6 flirt tool (Jenkinson and Smith, 2001; Jenkinson et al., 2002) with a normalised mutual information cost function, six degrees of freedom, and spline interpolation. Second, the resulting transformation matrices were applied to the ses-01 and ses-03 datasets, which were then linearly transformed and concatenated with ses-02 and ses-04, respectively. Finally, following concatenations, the datasets were normalised to mitigate remaining inter-session differences in signal intensity. Specifically, the non-diffusion-weighted MR volumes associated with each of the four sessions were averaged, and then the diffusion-weighted MR volumes from each session were divided by the corresponding averaged non-diffusion-weighted volumes, thereby generating the test and retest datasets. Hence, the datasets from test and retest variants represents normalised diffusion-weighted MR signal for each gradient direction ***g_i_***, i.e., *E* (***g****_i_*)=*S* (***g****_i_*)/ *S* (0), where *S* (***g****_i_*) is the diffusion-weighted MR signal for gradient direction ***g_i_*** and *S* (0) is the corresponding averaged non-diffusion-weighted signal across all non-diffusion-weighted volumes withing the session. These data are considered preprocessed. The preprocessed data use a single-precision floating-point (float32) data type.

### 2.4. Quantitative and qualitative modelling of the brain tissue

#### Microstructural measures

Microstructural measures: To demonstrate the practical feasibility of the dataset, we estimated a range of microstructural measures using four signal representations (IVIM, DTI, DKI, MAP-MRI), one biophysical model (NODDI) and one signal representation enabling the extraction of compartment-specific features (SMT). The techniques employed for technical validation use different experimental setups, including different numbers of shells and *b*-value configurations. The experimental setups were as follows:

- IVIM (Le Bihan, 2019): *b* = {0, 10, 20, 30, 50, 80, 100, 200, 300, 400, 500, 1000} s/mm^2^. The IVIM parameters were estimated using the variable projection-based algorithm (Farooq et al., 2016) implemented in DIPY library v1.12.0 (VarPro; dipy.reconst.ivim). We computed the following measures: perfusion volume fraction (*f _p_*), diffusion coefficient (*D*) and a pseudo-diffusion constant (*D*\*).
- DTI (Basser et al., 1994b): *b* = {0, 500, 1000} s/mm^2^. We considered two variants: 1) unconstrained ordinary least squares fitting implemented in FSL v6 (OLS; dtifit) that might produce negative eigenvalues, which form a physically implausible representation, and 2) an in-house implementation of the constrained non-linear Cholesky decomposition based algorithm (NLLS-Cholesky; Koay et al., 2006) that assumes the diffusion tensor is positively defined (i.e., all eigenvalues are strictly positive). We computed the following measures: fractional anisotropy (FA), mean diffusivity (MD), axial diffusivity (AD) and radial diffusivity (RD).
- DKI (Jensen et al., 2005): *b* = {0, 1000, 1400, 1800, 2200} s/mm^2^. We considered two variants: 1) unconstrained optimisation using the weighted least squares procedure implemented in DIPY v1.12.0 (WLS; dipy.reconst.dki), which may provide a physically implausible result, and 2) a constrained WLS (CWLS; `dipy.reconst.dki`) to enforce the non-negativity of the representation. We computed the standard DKI measures: mean kurtosis (MK), axial kurtosis (AK), radial kurtosis (RK), and three kurtosis tensor-based measures: mean kurtosis tensor (MKT), radial kurtosis tensor (RKT) and kurtosis fractional anisotropy (KFA).
- MAP-MRI (Özarslan et al., 2012): *b* = {0, 500, 1000, 1400, 1800, 2600, 3000} s/mm^2^. The MAP-MRI parameters were estimated using the DIPY library v1.12.0 (dipy.reconst.mapmri). The propagators were estimated using two variants: 1) unconstrained non-regularised optimisation (Özarslan et al., 2012), which may yield negative values of the propagator and therefore physically implausible solutions, and 2) a positivity constrained optimisation that enforces a non-negative propagator (Haije et al., 2020). In both cases, we used a maximal radial order of *L*_max_={4, 6 } and computed the following measures: the return-to-the-origin probability (RTOP), the return-to-the-axis probability (RTAP), the return-to-the-plane probability (RTPP), the mean-squared displacement (MSD), and the q-space inverse variance (QIV).
- NODDI (Zhang et al., 2012): *b* = {0, 700, 1000, 1800, 2200, 2600} s/mm^2^. The parameters were estimated using the Accelerated Microstructure Imaging via Convex Optimization (AMICO) framework v2.1.1 (Daducci et al., 2015). We computed the following measures: orientation dispersion (OD), intra-cellular volume fraction (*v*_ic_) and isotropic volume fraction (*v*_iso_).
- SMT (Kaden et al., 2016): *b* = {0, 500, 1000, 1400, 1800} s/mm^2^. Two parameters were estimated from the multi-shell SM-transformed signal, namely the free-water volume fraction (FWVF) and the diffusivity perpendicular to the axon *D*_perp_ (Tristán-Vega et al., 2022) using the dMRI-Lab toolbox (`atti2micro`; Tristán-Vega et al., 2025). The parallel diffusivity was fixed at *D*_par_=2.1×10^−3^ mm^2^/ s in the optimisation procedure.

All available normalised non-diffusion-weighted volumes were used in the above-mentioned methods to improve the robustness of the numerical procedures.

### FA registration to the common space

To retrieve brain white matter masks and assess the reproducibility of the microstructural measures, the DTI-based FA from the test and retest datasets were warped to the Montreal Neurological Institute (MNI) space, hereafter referred to as the common space. First, the FA maps computed from the *b* = {0, 500, 1000} s/mm^2^ were linearly registered to the `FSL_HCP1065_FA_1mm` template in the common space using the FSL v6 flirt tool (seven degrees of freedom, a normalised correlation cost function and spline-based interpolation; Jenkinson and Smith, 2001; Jenkinson et al., 2002). Second, the resulting affine transformations were refined using nonlinear registration with FSL v6 fnirt. Finally, the inverse transformations from the common space to each subject’s space were estimated using the FSL `invwarp` tool and used to warp the white matter labels from the Johns Hopkins University (JHU) atlas (Mori et al., 2025) to the corresponding native spaces using the FSL `applywarp` with the nearest-neighbour interpolation (`--interp=nn`). All microstructural measures were subsequently warped to the common space using the `applywarp` tool with a trilinear interpolation (`--interp=trilinear`) for the reproducibility analysis. We also computed the brain masks in native subjects’ spaces using the FSL `bet2` tool.

#### Reproducibility analysis of microstructural measures

We adopted the reproducibility analysis scheme presented in Pieciak et al., (2026b) for all microstructural measures estimated from the dataset. The spatially-varying variability coefficient has been defined in the common space for each measure as follows:

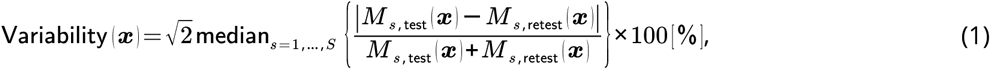

where *S* =11 is the number of subjects, and *M_s_*_,test_ (***x***) and *M_s_*_,retest_ (***x***) are the values of the measure from test and retest acquisitions in the common space at spatial position ***x*** for *s*–th subject. To quantitatively assess the reproducibility over the white matter, we used the labels from the JHU atlas. The reproducibility analysis was carried using an in-house script written in Python programming language and NumPy library.

#### Tractography

Biologically informed whole-brain tractograms were generated using the MRtrix3 software (Tournier et al., 2019) and white matter bundles were identified and registered using the DIPY library v1.12.0. First, fibre orientation distribution functions were estimated using the multi-shell multi-tissue constrained spherical deconvolution (Jeurissen et al., 2014) at *b* = {0, 500, 1000, 1400, 1800, 2600, 3000} s/mm^2^, followed by intensity normalisation. Second, anatomically constrained tractography (Smith et al., 2012), seeded at the grey matter-white matter interface (`5tt2gmwmi`, MRtrix3), was run to generate 5 million streamlines per brain (`tckgen`, MRtrix3). These streamlines were then filtered (i.e., reduced to 1.5 million streamlines per brain) using the SIFT algorithm with the MRtrix3 `tcksift` tool (Smith et al., 2013). Finally, the streamlines were linearly registered to the HCP-842 tractography atlas (`dipy.align.streamlinear.whole_brain_slr`; Garyfallidis et al., 2015; Yeh et al., 2018), and three individual white matter bundles were identified using the RecoBundles algorithm (`dipy.segment.bundles.RecoBundles`; Garyfallidis et al., 2018), namely the corticospinal tract (CST), arcuate fasciculus (AF), andinferior longitudinal fasciculus (ILF). To quantify the geometric similarity between the corresponding test and retest white matter bundles, we corrected the streamlines for spatial misalignment by linearly registering the test streamlines to the corresponding retest streamlines (`dipy.align.streamline_registration`; Garyfallidis et al., 2015), and then computed the weighted Dice coefficient (wDice), as defined by Cousineau et al. (2017)

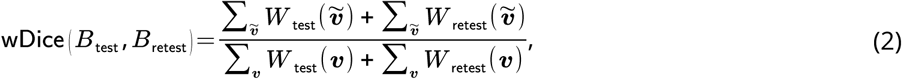

where *B*_test_ and *B*_retest_ are the white matter bundles reconstructed from the test and retest acquisitions with *W* _test_ (***v***) and *W* _retest_ (***v***) being the corresponding normalised fractions of streamlines passing though the voxel ***v*** defined in the spatial domain of the retest bundle, and ^∼^***v*** is the voxel within the logical intersection of the bundles *B*_test_ and *B*_retest_. Compared to the standard Dice coefficient, which measures only the binary spatial overlap, the wDice coefficient gives more importance to the voxels with higher streamline densities.

## 3. Data records

The dataset has been organised according to the BIDS format (Brain Imaging Data Structure; Gorgolewski et al., 2016) and is available in the OpenNeuro repository under the following link: https://openneuro.org/datasets/ds008695. The structure of the dataset has been validated to be BIDS compliant using the BIDS Validator (https://github.com/bids-standard/bids-validator). The size of raw dataset is 8.3GB, while the total size of preprocessed dataset is 39.5GB. The difference in size is due to the numerical representation of the data, i.e., the original data are stored as int16, whereas the preprocessed data are stored as float32. The general file tree structure is shown in Fig. 2.

**Figure 2.**
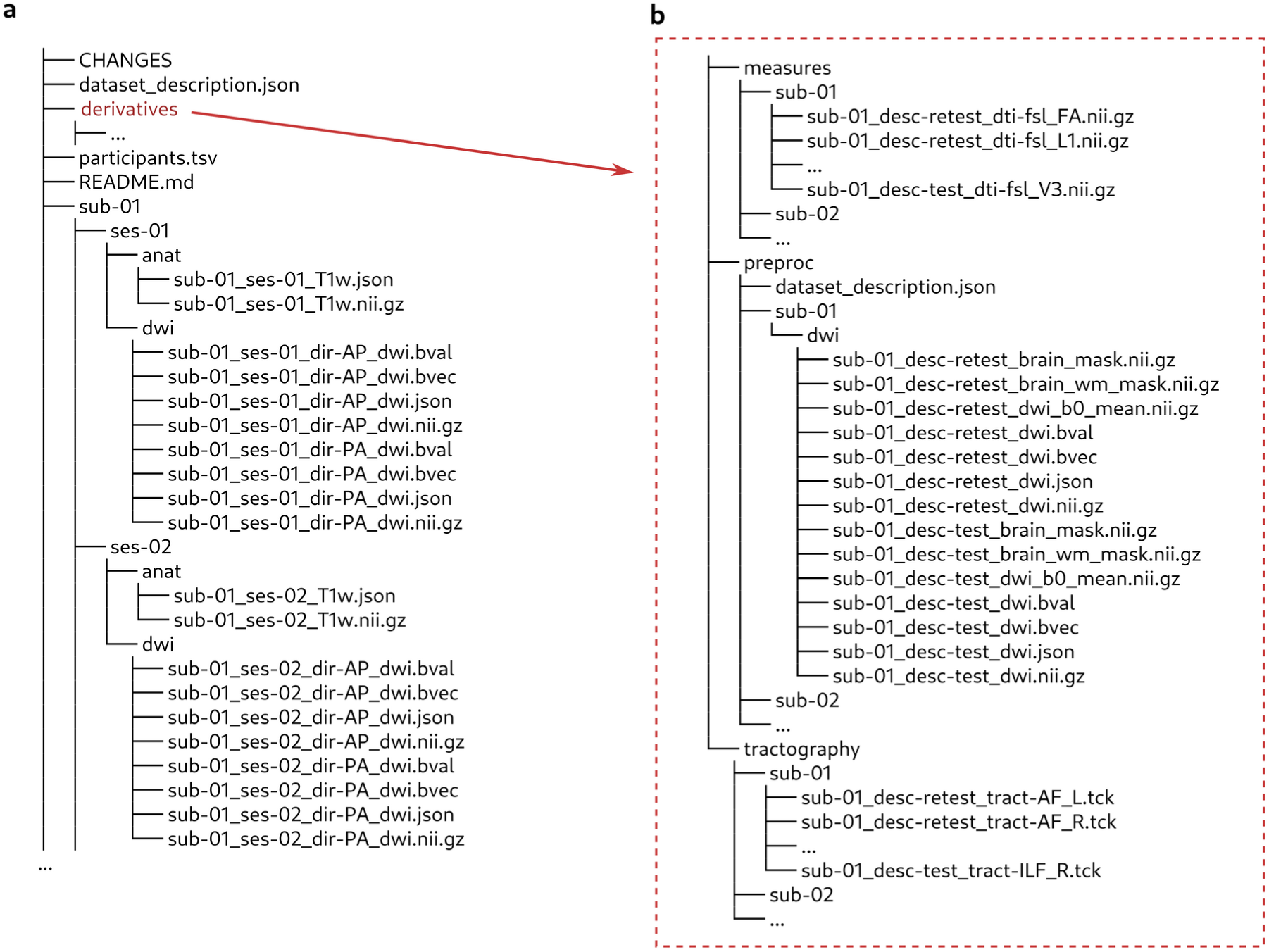
File tree structure of the dataset: **a)** general organisation of the raw data and **b)** organisation of the derived data, including preprocessed data (`/derivatives/preproc/`), DTI parameters estimated using the FSL software (`/derivatives/measures/`), and tractography results (`/derivatives/tractography/`).

### Raw data

Each volunteer’s raw data is provided in a separate folder entitled sub-XY (XY ∈ {01, 02, …, 11}), divided into four subfolders related to four acquisition sessions, as illustrated in Fig. 2a. For each session, ses-0Z (Z ∈ {1, 2, 3, 4}), face-removed *T_1_*-weighted NIfTI archive and JSON file are available in the folder /sub-XY/ses-0Z/anat/, while the diffusion-weighted MR data along with the *b*-values, gradient directions and JSON files are provided in the folder /sub-XY/ses-0Z/dwi/. Diffusion-weighted and non-diffusion-weighted volumes acquired in the AP encoding direction were coded as sub-XY_ses-0Z_dir-AP_dwi.nii.gz, while the corresponding non-diffusion-weighted volume acquired in the PA encoding direction is coded as sub-XY_ses-0Z_dir-PA.nii.gz. All raw volumes are NIfTI archives compressed using gzip.

### Preprocessed data

Fully preprocessed datasets are provided in a separate folder entitled /derivatives/preproc/. The folder consists of fully preprocessed and concatenated datasets, i.e., sessions ses-01 and ses-02 were merged into test datasets sub-XY_desc-test_dwi.nii.gz, while sessions ses-03 and ses-04 were merged into retest datasets sub-XY_desc-retest_dwi.nii.gz (see Fig. 2b). The folder also includes binary masks of the brain `sub-XY_desc-VARIANT_brain_mask.nii.gz (VARIANT` ∈ `{test, retest})`, white matter masks obtained from the JHU atlas `sub-XY_desc-VARIANT_brain_wm_mask.nii.gz`, and the averaged non-diffusion-weighted volumes `sub-XY_desc-VARIANT_dwi_b0_mean.nii.gz`. The *b*0 data were obtained by averaging all non-diffusion-weighted volumes from sessions ses-02 (test data) and ses-04 (retest data). All preprocessed volumes are NIfTI files compressed using gzip.

### Other derivative data

The folder /derivatives/ consists of two additional folders: /derivatives/measures/ and /derivatives/tractography/ (see Fig. 2b). The former holds DTI parameters estimated for test and retest data using the FSL dtifit tool, including the FA, eigenvalues (coded as L1, L2 and L3) and eigenvectors (coded as V1, V2 and V3). Each file has the following form: `sub-XY_desc-VARIANT_dti-fsl_MEASURE.nii.gz`, where `MEASURE` ∈ {FA, L1, L2, L3, V1, V2, V3}. The latter folder consists of fibre bundles reconstructed for test and retest data for both hemispheres of the brain. All bundles have been coded as: `sub-XY_desc- VARIANT_tract-TRACT.tck` with `TRACT` defined as `TRACT` ∈ {AF_L, AF_R, CST_L, CSR_R, ILF_L, IFL_R}.

### Other files

File `CHANGES` represents dataset version history. File `data_description.json` provides general information of the dataset coded in the JSON format. File participants.tsv presents tabularized meta-information for each subject, i.e., sex, age and handedness. File README.md codes webpage information about the dataset in Markdown markup language available under the link https://openneuro.org/datasets/ds008695. A separate file `data_description.json` is available for preprocessed dataset in the folder `/derivatives/preproc/`.

## 4. Technical validation

To assess the quality and demonstrate the utility of the dataset, we conducted several experiments, which can be generally grouped into three categories: 1) data quality, artefacts and preprocessing analysis (Figs. 3–5), 2) estimation of microstructural parameters (Figs. 6–8) and 3) reconstruction of fibre bundles (Fig. 9), with validation of the properties and reproducibility of both microstructural parameters and fibre bundles.

**Figure 3.**
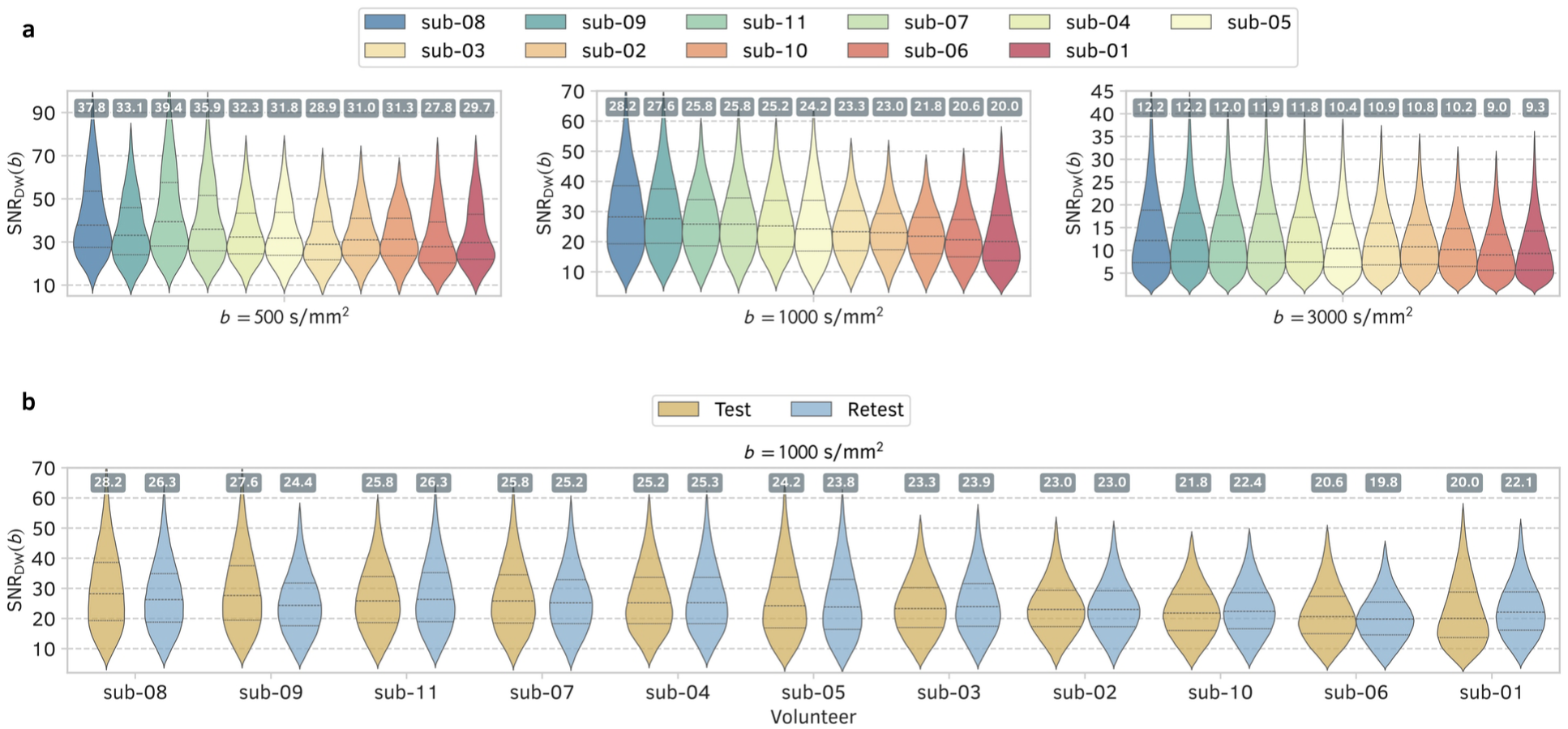
**a)** Diffusion-weighted signal-to-noise ratio SNR_DW_(*b*) computed for each volunteer at three selected *b-*values using test datasets. The numbers above the violin plots indicate the median SNR_DW_(*b*) within the white matter area. The plots are sorted in descending order according to the median SNR_DW_(*b*) at *b* = 1000 s/mm^2^. **b)** Test-retest SNR_DW_(*b*) for all volunteers evaluated at *b* = 1000 s/mm^2^. The plots are sorted in descending order according to median SNR_DW_(*b*) of the test datasets.

**Figure 4.**
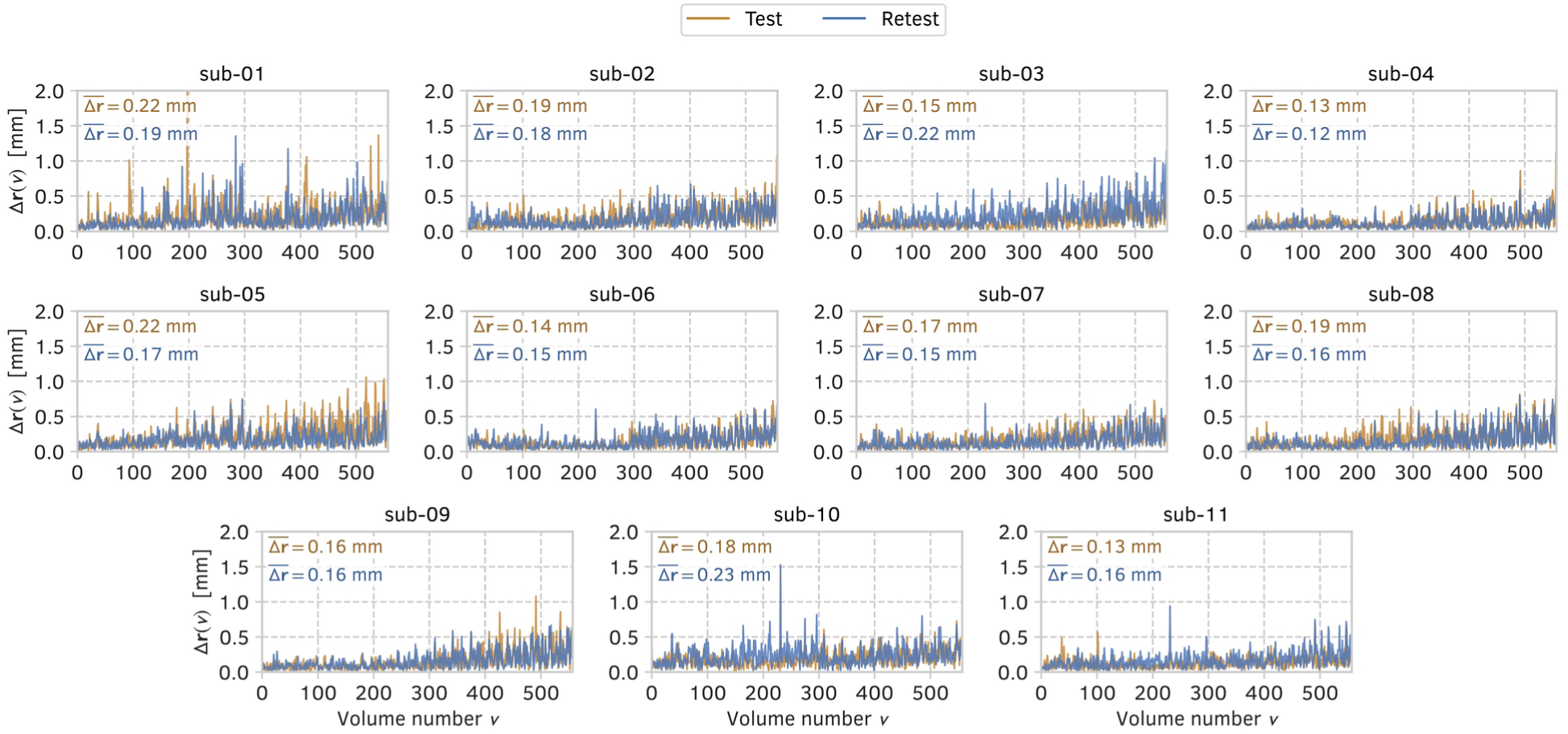
Head displacement, Δ**r** [*v*] (expressed in [mm]), across the volumes in the test and retest datasets. The index *v* corresponds to the volume index of the datasets in the acquisition protocol, as illustrated in **Fig. 1b**. The quantity 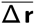 refers to the mean displacement across all volumes.

**Figure 5.**
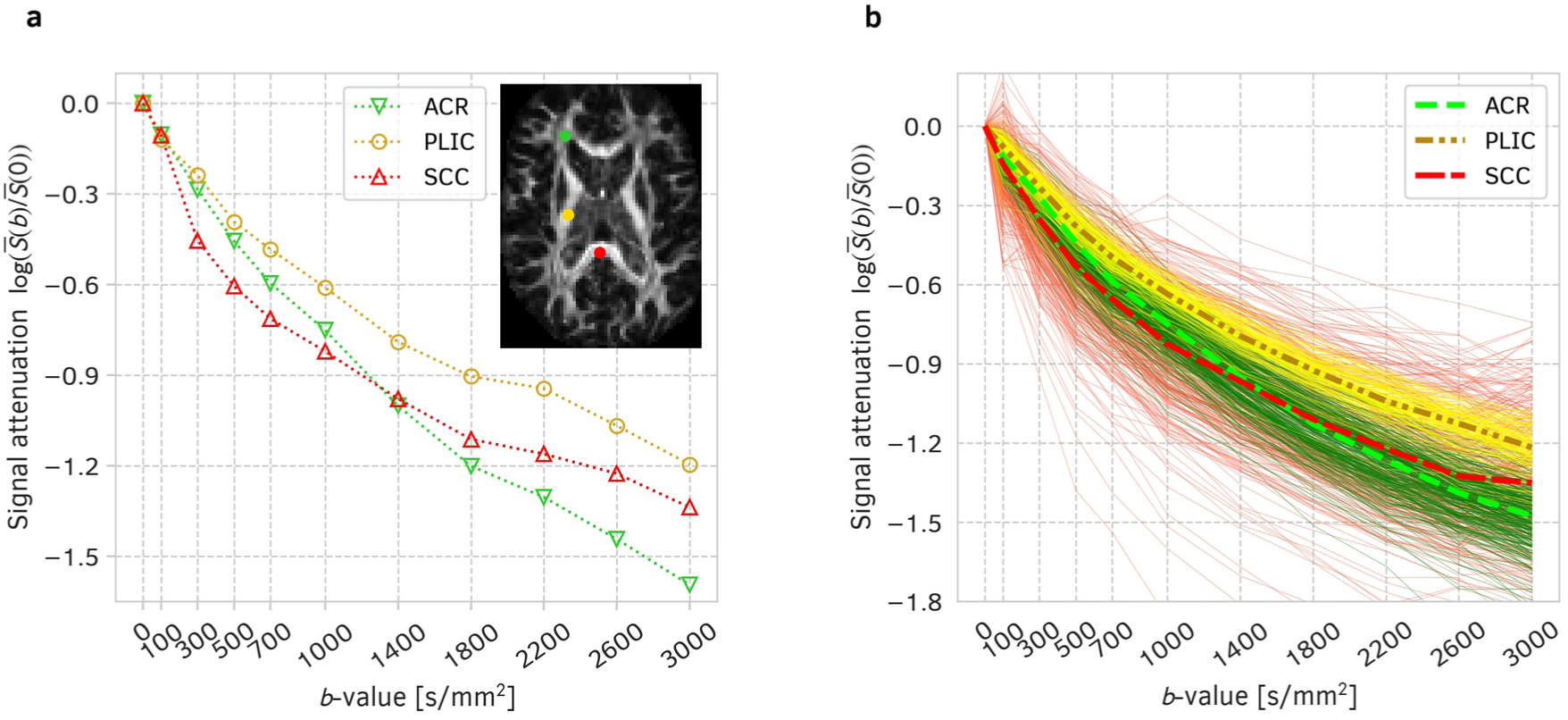
Data registration and concatenation results presented for test data of sub-02: **a)** log-transformed normalised spherical mean of diffusion-weighted MR signal as a function of *b*-value, log(*S* (*b*)/ *S* (0)), illustrated for three selected voxels taken from the anterior corona radiata (ACR), posterior limb of internal capsule (PLIC) and splenium of corpus callosum (SCC). **b)** Log-transformed normalised spherical mean signals obtained from the whole ACR, PLIC and SCC regions and presented as a function of the *b*-value. Each thin line represents the spherical mean signal at a single voxel with colors corresponding to those in panel a), while the thick dashed lines indicate the representatives (medians) computed over the region.

**Figure 6.**
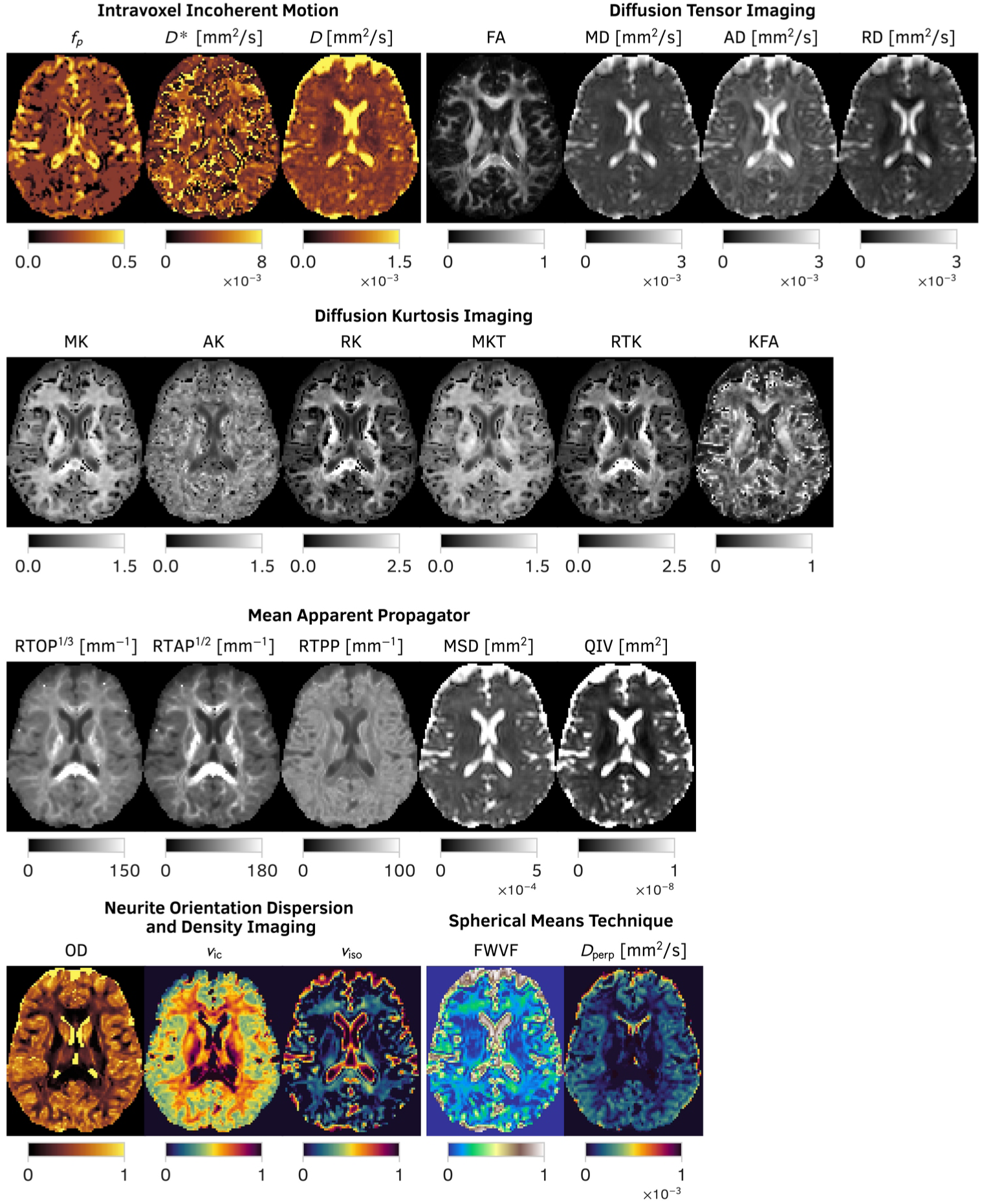
Visual inspection of microstructural measures obtained from a single subject (sub-04, test data) using the Intravoxel Incoherent Motion (IVIM), Diffusion Tensor Imaging, Diffusion Kurtosis Imaging, and Mean Apparent Propagator MRI representations, as well as the Neurite Orientation Dispersion and Density Imaging (NODDI) biophysical model and the Spherical Mean Technique (SMT). For visualisation purposes, non-grayscale colourmaps were applied to the IVIM, NODDI and SMT-based metrics.

**Figure 7.**
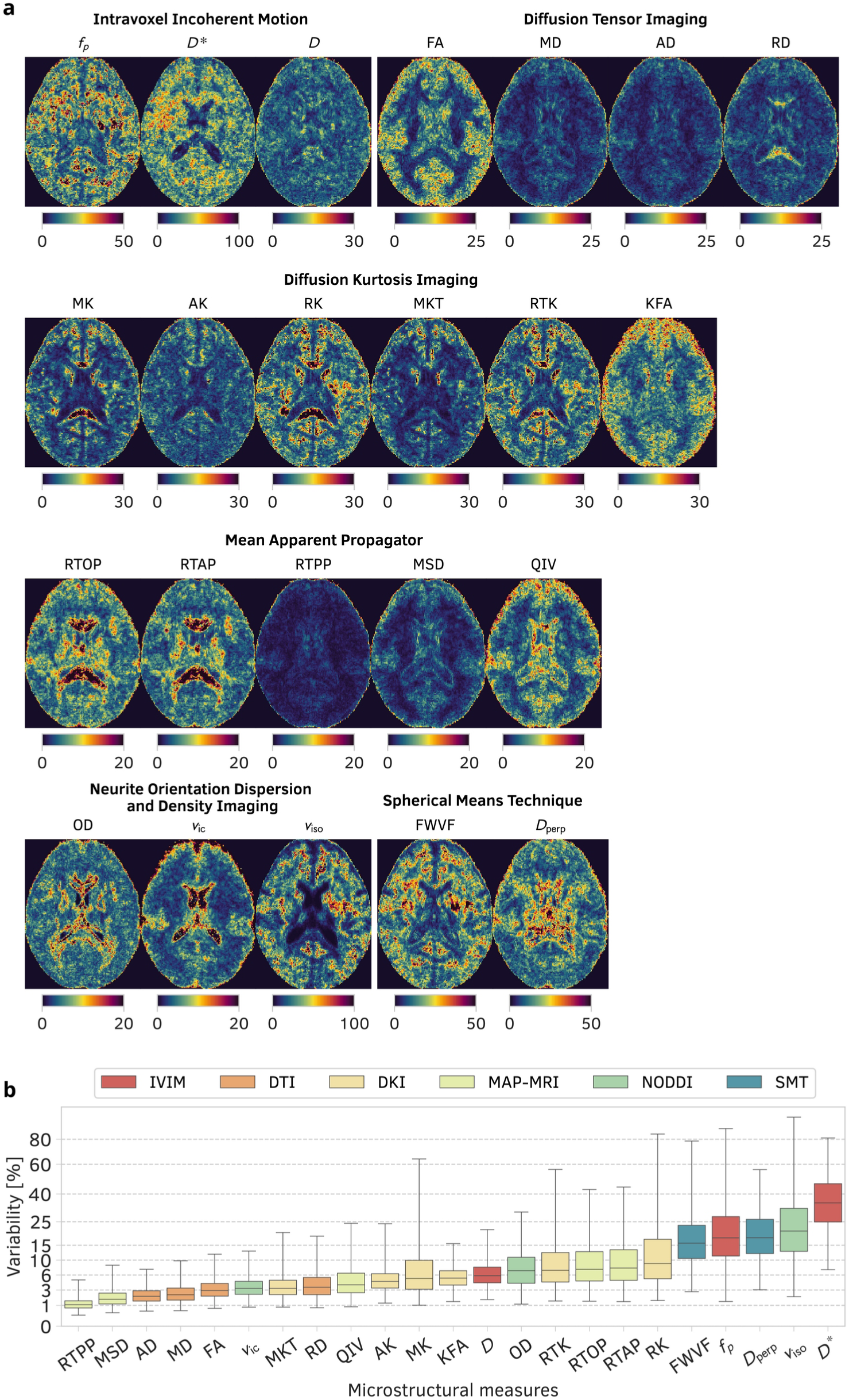
**a)** Spatially varying variability maps defined in the common space computed according to Eq. (1) for all microstructural parameters previously depicted in Fig. 6. The variability maps are expressed as percentages [%]. **b)** Boxplots illustrating the variability across the white matter region for all microstructural measures considered in this study. Each boxplot shows the first quartile (*Q*_1_), second quartile (median), and third quartile (*Q*_3_) of the variability index over the white matter from the common space. The whiskers indicate the range of observations within [*q*_0.005_, *q*_0.995_]. See the "Reproducibility analysis of microstructural measures" paragraph in section 2.4 for more details on the computation of the variability parameter.

**Figure 8.**
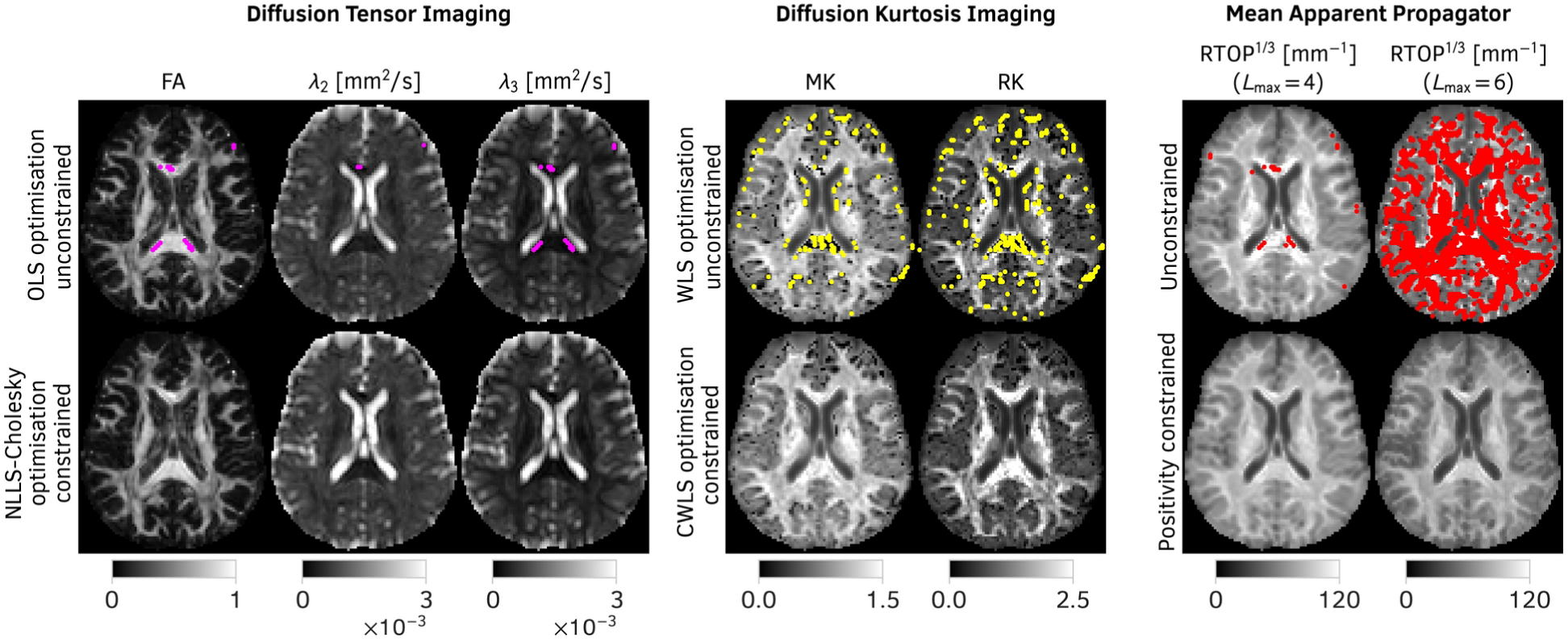
Microstructural measures estimated from sub-01 test data using unconstrained optimisation (top rows) and constrained optimisation procedures (bottom rows) from three different signal representations: Diffusion Tensor Imaging (DTI), Diffusion Kurtosis Imaging (DKI) and Mean Apparent Propagator MRI (MAP-MRI). Magenta, yellow and red voxels indicate implausible estimates due to the violation of physical constraints. Points marked in magenta refer to implausible DTI results (i.e., the second eigenvalue *λ*_2_ and/or third eigenvalue *λ*_3_ of the diffusion tensor are negative), yellow points present implausible DKI results (i.e., the value of MK and RK is lower than -3/7), while red points denote negative propagators in the MAP-MRI technique in at least one evaluated direction. The RTOP measure has been non-linearly scaled using a function *f* (*x*)=*x*^1/3^ for visualisation purposes and is given in [mm^−1^]. The parameter *L*_max_ denotes the maximal radial order of the MAP-MRI representation.

**Figure 9.**
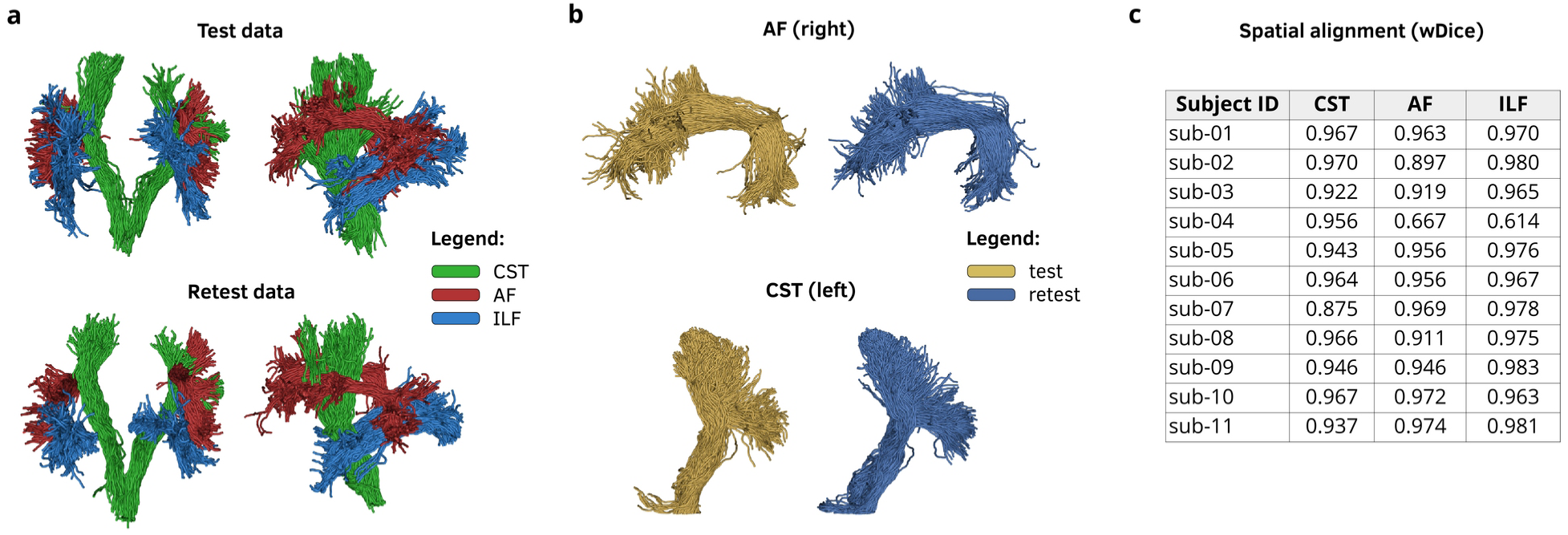
**a)** Fibre tracking results for sub-05 (test and retest datasets) illustrating three white matter bundles: the corticospinal tract (CST), arcuate fasciculus (AF), and inferior longitudinal fasciculus (ILF). **b)** Visual inspection of the spatial alignment between the test and retest right AF and left CST bundles (sub-05). **c)** The weighted Dice (wDice) coefficient was computed according to Eq. (2) for all subjects and presented as the mean of the left and right bundles.

### Signal-to-noise ratio

Fig. 3 shows the violin plots illustrating the diffusion-weighted signal-to-noise ratio aggregated over the white matter region derived from the JHU atlas. The SNR has been defined at each spatial location ***x*** as the ratio of denoised magnitude data to the spatially variant noise map estimated with the MP-PCA method. The SNRs from all spatial locations ***x*** at a specific *b-*value are aggregated and form a single violin plot defined as SNR_DW_(*b*). Before computing the violin plots, however, we eroded the white matter region with a structuring element of size 2 × 2 × 2 and additionally removed all SNR values that were outside the interval [*q*_0.005_, *q*_0.995_] with *q*_0.005_ being the quantile at 0.005. Fig. 3a presents the violin plots for the SNR_DW_(*b*) at *b* = 500, 1000, and 3000 s/mm^2^, ordered in descending order according to the median value computed across the white matter area of the test data at *b* = 1000 s/mm^2^. This experiment shows that 1) the differences in the SNR_DW_(*b*) across the subjects are more visible at lower *b*-value regime than at higher *b*-values, and 2) the highest SNRs are observed for sub-08, sub-09 and sub-11, while the lowest SNRs are observed for sub-01. Fig. 3b compares the SNR_DW_(*b*) computed for test and retest data at *b* = 1000 s/mm^2^ across all volunteers, ordered in a descending order according to the median value of the test data. In general, we observe that the dataset exhibits comparable SNRs across test and retest scans.

### Head displacements

Fig. 4 presents head displacements across the volumes in the test and retest datasets defined as 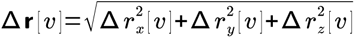 for *v*–th volume. For each test and retest dataset, we computed the mean head displacement 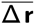 (defined in [mm]). The mean head displacement across all volumes was between 0.12 mm (sub-04, retest data) and 0.23 mm (sub-10, retest data). In general, the head displacements in all subjects were small, with the smallest observed in sub-04 and sub-06. The maximal displacement between two consecutive volumes was 2.1 mm for sub-01. No other significant head movements were detected in the cohort. We note that these head displacements were compensated using the FSL `eddy` tool.

### Data concatenation validation

In the next experiment presented in Fig. 5, we demonstrate data registration and concatenation results between sessions, i.e., ses-01 → ses-02 and ses-03 → ses-04. Particularly, we illustrate how the fully preprocessed diffusion-weighted MR data change as a function of *b*-value. Fig. 5a depicts the log-transformed normalised spherical mean of diffusion-weighted MR data, log(*S* (*b*)/ *S* (0)), for three selected voxels from the white matter of sub-02 test data, namely the anterior corona radiata (ACR), posterior limb of internal capsule (PLIC) and splenium of corpus callosum (SCC). The spherical mean signal has been computed as the average across all gradient directions available per *b*-value. Fig. 5b demonstrates the aggregated voxels from the ACR, PLIC and SCC regions (a single thin line represents the spherical mean from a single voxel as a function of *b-*value) and their representatives, computed as the median value across all voxels within the respective regions. This experiment confirms that the test and retest datasets were properly registered and merged across sessions. Additionally, we note that a slight noise floor bias can be observed for a higher *b*-value regime, which we decided not to correct.

### Visual inspection of microstructural measures

We now move on to experiments demonstrating the microstructural measures computed from the IVIM, DTI, DKI and MAP-MRI signal representations, the NODDI biophysical model and the multi-shell SMT. We used the experimental setups presented in section 2.4. In the case of DTI, DKI and MAP-MRI signal representations, we estimated the propagators using standard unconstrained optimisation procedures. MAP-MRI was estimated using a maximal radial order of *L*_max_=4. The results of this experiment for sub-04 test data are depicted in Fig. 6. We note that the DKI measures reveal ragged regions in white matter tissue and at tissue boundaries (e.g., white matter–cerebrospinal fluid), which are physically implausible and result from the unconstrained optimisation procedure. This effect is studied in more detail in the experiment depicted in Fig. 8. All in all, as the experiment demonstrates, one can obtain a broad spectrum of microstructural measures using various mathematical models to characterise the diffusion-weighted MR signal.

### Reproducibility analysis

Next, in Fig. 7, we present the results of the reproducibility analysis for all measures previously presented in Fig. 6. Specifically, we first transformed all measures estimated for all subjects (test and retest data) to the common space and then computed the spatially-varying variability coefficient for each measure, as defined in Eq. (1). The results presented in Fig. 7a illustrate the median variability maps (represented by the coefficient of variation defined in %) of the measures across all subjects in the dataset. In Fig. 7b, we quantify these variabilities by presenting boxplots for each measure over the white matter area. In general, we can observe that the DTI- and MAP-MRI- based measures are characterised by the smallest variability (i.e., reproducibility is the highest).

### Unconstrained vs constrained optimisation

In the next experiment, we depict microstructural measures computed from DTI, DKI and MAP-MRI representations using unconstrained and constrained optimisation techniques. The propagator in the MAP-MRI technique was obtained from its regularised version with maximum radial orders of *L*_max_=4 and *L*_max_=6. The assumption behind this experiment is to demonstrate how an unconstrained optimisation may yield physically implausible results. In Fig. 8, we present estimated microstructural maps for sub-01 test data with points marked in magenta, yellow and red, where the optimisation resulted in physically implausible results. For example, in DTI a pixel marked in magenta indicates that the second eigenvalue *λ*_2_ and/or third eigenvalue *λ*_3_ of the estimated diffusion tensor is negative. In DKI, the pixel marked in yellow indicates that the value of MK or RK is lower than -3/7, which is the theoretical lower bound of kurtosis for a region that consists of water confined to spherical pores (Jensen et al., 2005). In the case of MAP-MRI, red pixels denote negative propagators in at least one evaluated direction. All in all, this experiment emphasizes that the presented dataset is useful for verifying physical plausibility of the estimated propagators.

### Tractography

In the final experiment, we present fibre-tracking results. Specifically, we obtained whole-brain tractograms for test and retest data for all subjects and then identified three white matter bundles, as described in section 2.4. Fig. 9a,b illustrate these fibre bundles for sub-05, while Fig. 9c presents the quantitative evaluation of spatial alignment between test and retest bundles for all subjects using the wDice coefficient. Here, we observe high consistent spatial agreement across all subjects except sub-04. This subject has proven difficult for fibre bundle estimation, which highlights the utility of our dataset for supporting the development of new robust tractography algorithms.

## 5. Code availability

We used publicly available software packages to convert, preprocess and register the dataset, estimate microstructural parameters, and reconstruct fiber bundles. All processing was handled using Bash scripting under GNU/Linux and the Python programming language v3.13.7 with the NumPy v2.3.3 library.

## Data conversion

Data were converted from DICOM to NIfTI format using the dicm2nii tool v2026.03.19 (https://github.com/xiangruili/dicm2nii). The JSON files were generated using dicm2nii and then pseudonymised and corrected using in-house software written in Python v3.13.7. Face masking of T1-weighted data was performed using the FSL FMRIB Software Library v6 (https://fsl.fmrib.ox.ac.uk/).

## Preprocessing

Noise estimation, denoising and Gibbs ringing artefacts correction were performed using MRtrix3 (https://www.mrtrix.org/). Skull stripping, correction of susceptibility-induced distortions and eddy-current artefacts, and data registration between sessions and to MNI space were performed using the FSL FMRIB Software Library v6. B1 field inhomogeneity was corrected using ANTs v2.6.3 (https://github.com/antsx/ants). Time-varying drift was corrected using software provided by the authors of the method (https://es.mathworks.com/matlabcentral/fileexchange/55008-signal-drift-correction-for-diffusion-mri-data).

Microstructural parameters: Unconstrained DTI was estimated using FSL v6. Constrained DTI with the NLLS Cholesky method was estimated using in-house software written in Python programming language and run under Python v3.13.7 with NumPy v2.3.3 and SciPy v1.16.2. IVIM, DKI and MAP-MRI parameters were estimated using the DIPY library v1.12.0 (https://dipy.org), NODDI via AMICO v2.1.1 (https://github.com/daducci/AMICO) and SMT-based metrics using the dMRI-Lab toolbox (https://github.com/atriveg/dmrimatlab) run under MATLAB R2025b (The MathWorks, Inc., Natick, MA).

## Tractography

Tractography was performed using MRtrix3, registered between test and retest data using DIPY v1.12.0 and visualised with DIPY v1.12.0 and VTK v9.5.2 (https://vtk.org/). The wDice coefficient was computed using in-house software written in Python.

## Acknowledgements

This work was funded by the Agencia Estatal de Investigación (Ministerio de Ciencia, Innovación y Universidades of Spain) with the research grant PID2024-158963NB-I00. Irene Guadilla is supported with the Research Grant VA156P24 funded by Junta de Castilla y León (Consejería de Educación) and the European Regional Development Fund (ERDF/FEDER).

## Author contributions

T.P. initiated and conceptualized the project, collected, preprocessed and registered data, computed microstructural parameters, designed the experiments, generated all figures, wrote the original manuscript, T.P. and R. M.B. prepared the acquisition procedure, D.C. prepared tractography experiment code, T.P., I.G., R. N.-G., S. M.-C., P. V.-A., L. M.H., M. V.A, A. T.V. reviewed and edited the manuscript, A. T.V. and T.P. acquired funding. All authors approved the manuscript.

## Declaration of generative AI

The authors did not use generative AI in writing of the manuscript.

## Additional information

Correspondence and requests for materials should be addressed to T.P.

